# Correcting False Memories: The Effect of Mnemonic Generalization on Original Memory Traces

**DOI:** 10.1101/2020.04.13.039479

**Authors:** Nathan M. Muncy, C. Brock Kirwan

**Affiliations:** Department of Psychology, Brigham Young University, Provo, Utah, USA; Neuroscience Center, Brigham Young University, Provo, Utah, USA; Center for Children and Family, Florida International University, Miami, Florida, USA

**Keywords:** episodic memory, discrimination, generalization, fMRI

## Abstract

False memories are a common occurrence but the impact of misremembering on the original memory trace is ill-described. While the original memory may be overwritten, it is also possible for a second false memory to exist concurrently with the original, and if a false memory exists concurrently then recovery of the original information should be possible. This study investigates first, whether false recognition overwrites the original memory representation using a mnemonic discrimination task, and second, which neural processes are associated with recovering the original memory following a false memory. Thirty-five healthy, young adults performed multiple recognition memory tests, where the design of the experiment induced participants to make memory errors in the first recognition memory test and then allowed us to determine whether the memory error would be corrected in the second test. FMRI signal associated with the encoding and retrieval processes during the experiment were investigated in order to determine the important regions for false memory correction. We found that false memories do not overwrite the original trace in all instances, as recovery of the original information was possible. Critically, we determined that recovery of the original information was dependent on higher-order processes during the formation of the false memory during the first test, and not on processing at the time of encoding nor the second test.

## 1 Introduction

The field of study on mnemonic discrimination in episodic memory has largely focused on a single aspect—the capacity of the participant to correctly identify lure stimuli. In animal models, where pattern separation processes can be targeted, previous work has demonstrated that the successful identification of lures is dependent on the orthogonalization processes of the dentate gyrus in the hippocampus as well as an intact hippocampal trisynaptic pathway (Gilbert & Kesner, 2006; Gilbert et al., 1998; Gilbert et al., 2001; Gold & Kesner, 2005; Hunsaker et al., 2008; Kesner, 2007a, 2007b; Leutgeb et al., 2007). Human studies have implicated the same processes for successful mnemonic discrimination performance: participants with healthy hippocampi perform well on tests of mnemonic discrimination while participants with hippocampal atrophy perform poorly, and participants with hippocampal lesions perform worst of all (Bakker et al., 2008; Kirwan et al., 2012; Mueller et al., 2011; Small et al., 2011). Additionally, hippocampal CA3/DG volumes and activity are known to positively correlate with human performance on tests of mnemonic discrimination (Doxey & Kirwan, 2015; Grady & Ryan, 2017; Leal & Yassa, 2014; Marshall et al., 2016; Reagh et al., 2014; Riphagen et al., 2020; Roberts et al., 2014; Sheppard et al., 2016). That said, while mnemonic discrimination processes appear to be pattern separation dependent, higher order processes such as semantic labeling, visual spatial processing, attention, and decisiveness (Goebel & Vincze, 2007; Hunsaker & Kesner, 2013; Lacy et al., 2011; Leal & Yassa, 2014; Pidgeon & Morcom, 2016; Reagh et al., 2016; Reagh & Yassa, 2014; Rolls & Kesner, 2016; Steemers et al., 2016; Wais et al., 2017) integrate with hippocampal pattern separation output to coordinate the overall behavior. But while the roles of the medial temporal lobe structures in mnemonic discrimination are well understood, higher-order processes are relatively understudied in this context.

In addition to these gaps in the field of mnemonic discrimination, the failure in detecting lures is also relatively unaddressed. The detection of lure stimuli in a recognition memory test is the result of both encoding and retrieval processes: the lure’s representation must be successfully orthogonalized from the target’s, and the target must be successfully retrieved. At the point of successful lure identification, then, two distinct representations exist (Target, Lure) between which the participant is able to discriminate. This process is sometimes known as “recall to reject” (J. Kim & Yassa, 2013; Kirwan & Stark, 2007; Lacy et al., 2011; van den Honert et al., 2016). A failure in lure detection, then, could have multiple antecedents: a failure to encode the lure’s salient details, a failure to retrieve the salient target information, and/or lure interference all may lead to mnemonic generalization rather than discrimination. Such failures might result from under-performing lower-order hippocampal processes, but may also be the result of deficient attention, semantic labeling, and/or decision making. Further, and perhaps more interestingly, the consequence of incorrectly identifying the Lure as Target (a False Alarm, FA) is unknown: as studies of mnemonic discrimination typically use a single study/test presentation, the effect of the FA response on the original memory trace has not been established. It is possible that either the lure representation could overwrite the target memory trace, or that a second representation could exist concurrently with the original. If the latter is true and multiple memory traces result from a FA response, then the two memory traces (one of the Target, and one of the Lure) may be dissociable in subsequent testing. And while certain studies, particularly in the false-memory literature, have specifically targeted the FA responses (rather than record it as a task failure in the mnemonic discrimination task) and the underlying mechanism at both the encoding and test phase of the trial (Cabeza, 2013; Hanczakowski & Mazzoni, 2011; H. Kim, 2011; Kubota et al., 2006; Moritz et al., 2006; Okado & Stark, 2005), the differential processing that could potentially be involved with correcting versus perpetuating FA behavior has not been investigated.

In order to address these gaps in the literature, we developed a paradigm to investigate both the impact of a false alarm on the original memory trace as well as the relative contributions of hippocampal and cortical processes in promoting and recovering from a failed lure detection. We hypothesized that (a) FAs would result in multiple memory traces that exist concurrently, and as such, we predicted that participants would be able to distinguish target from lure when both were presented simultaneously. Further, we hypothesized that subsequent target-trace recovery would be predicted by (b) increased BOLD signal in CA3/DG during the orienting task and false alarm and (c) increased BOLD signal in *a priori* regions associated with encoding and retrieval. Finally, we performed exploratory whole-brain analyses with the prediction that (d) we would observe regions associated with higher-order cognitive processes, such as attention and content processing, that were involved in subsequent target-trace recovery. In short, we predicted that not all FA responses would be equal, but that some would occur during better processing of the Lure stimulus which would then precede a retrieval of the Target memory trace during a subsequent test.

## 2 Methods

### 2.1 Participants

Thirty-five participants (15 female, mean age = 23.1, SD = 2.2) were recruited from the university and local community and were compensated for their participation with either payment or a 3D-printed, quarter-scale model of their brain. The university’s Institutional Review Board approved the research, and all participants gave written informed consent prior to participation. Inclusion criteria consisted of speaking English as a native language, being right-hand dominant, and having normal or corrected-to-normal vision. Exclusion criteria consisted of a history of psychological disorder or brain injury, head dimensions that were too large for the 32-channel coil, the existence of non-surgical metal in the participant’s body (including non-removable piercings), and color blindness. Participants were excluded from MRI analyses separately for excess motion within a particular phase of the experiment as noted in Section 2.5.

### 2.2 Mnemonic Discrimination Task

Stimuli for the task consisted of 200 pairs of perceptually and semantically similar Target-Lure pairs that were randomly selected from those hosted at https://github.com/celstark/MST. Each stimulus depicted an everyday object in the center of the image with a white background, and one member of each set was randomly selected to be a Target while the other was selected to be a Lure for each participant. Previous to this study, approximately 900 Target-Lure pairs were rated for similarity by 35 individuals (who were independent of the current study) on a 7-point Likert scale where a response of “1” indicated “extremely dissimilar” and “7” indicated “extremely similar”. Only items that received an averaged similarity rating of 6.0 or higher were used in this task in order to ensure a sufficiently large number of FA responses for functional analyses.

The behavioral task consisted of three phases (Study, Test1, and Test2), all of which were performed while the participant was scanned using fMRI. Stimuli were displayed by means of an MR-compatible LCD monitor placed at the head of the scanner and viewed using a mirror mounted on the head coil. Responses were collected via a fiber optic button box. E-prime (v 2.0) was used to control stimulus display and record behavioral responses. While in the scanner but prior to engaging in the task, participants viewed a task instruction video; for consistency, the same researcher (NMM) screened and debriefed all participants, responded to any questions about the task, and operated the scanner. Three different instruction videos were shown, one immediately prior to each respective phase of the task, in which explicit directions and examples were given to the participant.

Study phase: In the initial encoding phase, participants were shown 200 images of everyday objects and were instructed to indicate via button press whether each object is typically encountered indoors or outdoors (Figure 1, left). Each stimulus was preceded by a fixation cross for 500 ms, then presented for 3500 ms across one continuous block. For all phases of the experiment, 16 blank trials were added in a pseudo-random fashion to create jitter in stimulus timing, with care taken to ensure that each block contained an equal number of blank trials (when applicable). These blank trials, in addition to the inter-stimulus fixation crosses, served as the baseline condition in the single subject regression analysis (see Section 2.5).

**Figure 1:**
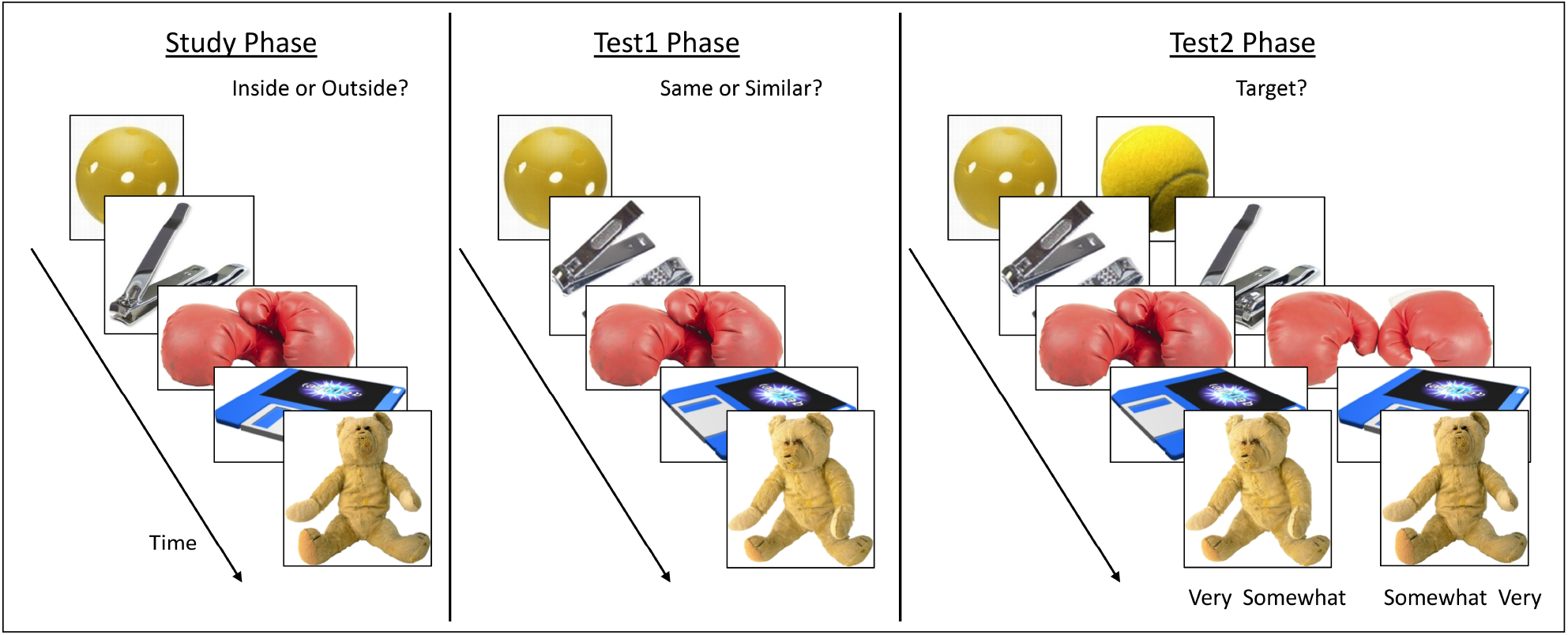
A representation of each phase of the behavioral task. Left, during the study phase the participants made indoor/outdoor judgments about a series of objects. Middle, during the first test phase, participants were presented with either the same (Target) stimulus or one that was similar (Lure) and were asked to decide whether the stimulus was the same or similar. Right, during the second test phase, participants were presented with both Target and Lure stimuli simultaneously and asked to indicate which item was originally presented during the Study phase. Additionally, participants indicated the level of confidence (Very or Somewhat) in their decision and made both their target and confidence decision with a single button press. For instance, if they decided that the left stimulus was the Target and they were very confident, they pressed “1” whereas if they were only somewhat confident the right stimulus was the Target, they pressed “3”.

Test1 phase: Participants were shown 100 Target stimuli (items identical to the original stimuli in the Study phase), and 100 Lure stimuli (items which were similar to, but differed visually from, the original items) one at a time in the center of the screen. Participants were instructed to indicate via button press whether each stimulus was the “same” (an exact repeat, or Target) or “different” (a related Lure) to the study item (Figure 1, middle). Stimuli were presented in a randomized order for both type (Target/Lure) and for level of Target-Lure similarity. The Test1 phase was conducted in two blocks of 100 stimuli each (50 Target, 50 Lure) and each stimulus was presented for 3500 ms and preceded by a fixation cross for 500 ms.

Test2 phase: In this phase, all stimuli from the Study phase (Targets), accompanied with the similar item (Lure), were presented side-by-side to the participant in a two-alternative forced-choice format. The stimuli were counterbalanced for whether the Target appeared on the Left or Right (Figure 1, right). Participants were asked to indicate which of the two items they had seen during the Study phase (Left or Right) and their confidence in their decision (Very or Somewhat) via one of four potential response options (left very confident; left somewhat confident; right somewhat confident; right very confident). This phase was conducted in four blocks of 50 stimuli in an attempt to combat participant fatigue. Each stimulus was presented for 5000 ms and preceded by a fixation cross for 500 ms. During all phases of this experiment, participants were instructed to respond while the stimulus was presented on the screen, and no feedback was given relating to participant performance.

### 2.3 Statistical Analyses of Behavioral Data

Participant responses during the Study phase of the experiment were not analyzed for indoor-outdoor accuracy as this question was merely meant to orient participants’ attention to the stimuli. During the second phase of the experiment, Test1, participants saw either Target or Lure stimuli and made a forced-dichotomous judgment of whether the item was “Same” or “Different”. Correctly identifying the Target and Lure is termed Hit and Correct Rejection (CR), respectively, and the incorrect identification is termed Miss and False Alarm (FA), respectively. The third phase of the experiment, Test2, was a two-alternative forced-choice where both Target and Lure stimuli were presented and the participant was asked to identify the Target; consequently, only Hits and FAs were possible.

Behavioral analyses were conducted on response proportions, considering the number of items to which each participant responded. D-prime (d’) scores were calculated for both Test1 and Test2 phases as an indicator of each participant’s sensitivity to the stimuli. Test1 d’ scores were calculated the standard way (Equation 1; Wickens, 2002) where Z = z-score, pHit = proportion of Hits to the number of Targets, and pFA = proportion of FAs to the number of Lures. As the Test2 phase presented both stimuli simultaneously, a correction was added to the d’ calculation (Equation 2; Wickelgren, 1968).

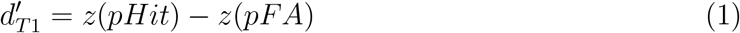

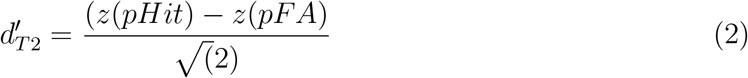

Further, one-sample proportion testing using a continuity correction were used to assess whether participants were more likely to respond one way versus another given a stimulus (e.g., if participants were more likely to Hit or Miss when presented with a Target, or more likely to respond Very versus Somewhat confident during CRs). Outliers were detected according to the 1.5 × IQR method. Paired and one-sample t-testing was used to compare the d’ scores to each other and against 0, respectively, utilizing an FDR correction to account for multiple comparisons (reported as *q*-values) when appropriate. When the exclusion of outliers differentially impacted the statistics, tests including and excluding outliers are reported, otherwise reporting represents the exclusion of outliers.

### 2.4 MRI Acquisition

Functional and structural images were acquired on a Siemens TIM Trio 3T MRI scanner utilizing a 32-channel head coil. Participants contributed a T1-weighted scan, a high-resolution T2-weighted scan, and a series of functional T2*-weighted scans as described below. Standard-resolution structural images were acquired using a T1-weighted magnetization-prepared rapid acquisition with gradient echo (MP-RAGE) sequence with the following parameters: TR = 1900 ms, TE = 2.26 ms, flip angle = 9°, FoV = 218 × 250, voxel size = 0.97 × 0.97 × 1 mm, slices = 176 interleaved. High-resolution structural images were acquired using a T2-weighted sequence with the following parameters: TR = 8020 ms, TE = 80 ms, flip angle = 150°, FoV = 150 × 150 mm, voxel size = 0.4 × 0.4 × 2 mm, slices = 30 interleaved. The T2 data were acquired perpendicular to the long axis of the hip-pocampus. High-resolution multi-band echo-planar image (EPI) scans were acquired with a T2*-weighted pulse sequence with the following parameters: Multi-band factor = 8, TR = 875 ms, TE = 43.6 ms, flip angle = 55°, FoV = 180 × 180 mm, voxel size = 1.8 mm^3^, slices = 72 interleaved. EPI scan slices were aligned parallel with the long axis of the hippocampus, and the first 11 TRs were discarded in order to allow for T1 equilibration.

### 2.5 MRI Data Pre-Processing

The pre-processing and visualization of structural and functional MRI data were accomplished with the following software: dcm2niix (Li et al., 2016), Convert 3D Medical Image Processing Tool (c3d) and ITK-SNAP (Yushkevich et al., 2006), Advanced Normalization Tools software (ANTs; Avants et al., 2008; Avants et al., 2011)), Automatic Segmentation of Hippocampal Subfields (ASHS; Yushkevich et al., 2015), and Multi-image Analysis GUI (Mango; Lancaster et al., 2012).

T1-weighted structural files were converted into 3D NIfTI files, rotated into functional space (referencing the volume with the minimum calculated outlier voxels) using a rigid transformation with a Local Pearson Correlation cost function. The rotated scan was then skull-stripped and warped into MNI space via a non-linear diffeomorphic transformation.

High-resolution T2-weighted files were converted to 3D NIfTI files, and both T1w and T2w files, in native space, were used to segment hippocampal subregions via ASHS, referencing the UPenn atlas (Yushkevich et al., 2015). Hippocampal subregion masks from each participant were then used to construct template priors via Joint Label Fusion (JLF; Wang et al., 2013), and study-specific masks for the CA1 and the combined CA2/CA3/DG regions were constructed and resampled into the same dimensions as the functional data. Voxels associated with overlapping subregions were excluded to combat partial-voluming effects when down-sampling. The CA2, CA3, and DG masks were combined into a single mask due to functional resolution constraints (Supplemental Figure S1).

T2*-weighted DICOMs were first converted to 4D NIfTI files, and the volume (or TR) with the smallest percentage of outlier voxels was extracted to serve as a volume registration base. The calculations for moving each volume into the same space as the volume registration base were then performed, using cubic polynomial interpolations, and the T1-weighted structural file was also aligned with the volume registration base using a rigid transformation that utilized a Local Pearson Correlation cost function. This rotated structural scan was then used to produce normalization calculations via a non-linear transformation into MNI space. Movement of the functional data into template space was then accomplished by concatenating the volume registration calculations with the MNI transformation calculations, thereby moving all volumes from native space to MNI space via a single transformation, resulting in less partial-voluming and blurring of the functional data than if each transformation was applied separately. A series of masks were then generated in order to remove volumes with missing data as well as determine the intersection between functional and structural data, and the functional data were then scaled by the mean signal. Finally, centered (demeaned) motion files for six degrees of freedom were generated for each run of the experiment.

Behavioral data during each of the three phases (Study, Test1, and Test2) were coded according to behavioral responses during that same as well as subsequent phases of the experiment. Trials from the Study phase were coded according to subsequent Test1 performance (i.e., Study preceding Test1 activation): subsequent Hit, subsequent FA, subsequent Miss, or subsequent CR. Study-phase trials were also coded according to behavioral responses in Test1 that preceded a Test2 behavior. For example, a stimulus presented during Study might subsequently receive a FA in Test1 and a Hit in Test2 (subsequent FA-Hit). Coding Study trials in this way allows us to potentially localize differences during the encoding process that result in Test1 FAs that are perpetuated into Test2 (subsequent FA-FA) compared to FAs that are corrected in Test2 (subsequent FA-Hit). It should be noted, however, that a low number of certain behaviors (particularly Test1 Misses preceding Test2 behaviors) necessitated collapsing across CR and Miss response bins in the deconvolution phase in order to avoid producing vectors for some participants consisting entirely of zeros, which are incompatible with deconvolution. Such behavioral bins were not included in group-level analyses. Test1 trials were coded according to four outcomes (Hit, FA, CR, and Miss). Responses during Test1 that preceded Test2 behaviors produced 4 × 2 possible outcomes since Test2 had two possible behaviors (Hit, FA). Test2 responses that followed Test1 also produced 4 × 2 possible outcomes, and Test2 had two outcomes. Note that dividing all Test2 behavioral data according to confidence interval as well as response type resulted in bins with trial counts that were too small for meaningful modeling of the BOLD signal, and as such the confidence interval was unfortunately dropped from the functional analyses. Finally, trials where participants failed to respond were coded separately from each of the above.

Vectors coding for the above behavioral outcomes were used in single-subject regression models (Generalized Least Square) in which the canonical hemodynamic response was convolved with a boxcar function with a duration equal to stimulus presentation. Modeling included polynomial regressors accounting for scanner drift and scan run and regressors coding for movement. The baseline condition consisted of collapsing across inter-stimulus intervals and jittered blank stimuli. Volumes consisting of movement greater than 0.3° and/or more than 0.3 mm in any direction relative to the previous volume were discarded along with the previous volume. Each deconvolution was assessed for the number of volumes that necessitated removal and any participant with more than 10% of volumes dropped and 1.5 × IQR of volumes dropped were excluded from subsequent analyses: one subject was removed from the Study phase, two subjects were removed from the Test1 phase, and three subjects were removed from the Test2 phase. As such 34 participants were used to investigate Study performance, 33 for Test1, and 32 for Test2 performance.

Region of interest (ROI) specific masks were constructed to test *a priori* hypotheses. In addition to the hippocampal subregion masks described above, clusters associated with memory tasks were constructed by using the key words “Memory Encoding” and “Memory Retrieval” as search terms in the NeuroSynth algorithm, which conducts a meta-analysis of PubMed articles reporting the same key words and constructs a set of clusters derived from the reported coordinates for contrasts with specific labels. The search term “Memory Encoding” yielded 124 articles, while “Memory Retrieval” yielded 183. Association clusters (corrected for meta-analysis multiple comparisons at the p=.01 level) were downloaded and thresholded at k=80 for the encoding clusters, and k=100 for the retrieval clusters, which produced a set of binary masks that were then resampled into functional dimensions. The “encoding” set contained four clusters, consisting of the left hippocampus, right collateral sulcus, left inferotemporal gyrus, and right amygdala (Supplemental Figure S2). The “retrieval” set contained 10 clusters, consisting of the left retrosplenial, angular gyrus, hippocampus, medial PFC, temporal-parietal junction, dorsal medial PFC, and the right hippocampus, parietal-occipital sulcus, and posterior hippocampus (Supplemental Figure S3). NeuroSynth masks were used to extract region-of-interest (ROI) specific mean *β*-coefficients. Two-factor repeated measures ANOVAs were used to investigate an ROI × behavior interaction for hippocampal subregion and NeuroSynth masks, individually, where a Greenhouse-Geisser (GG) correction was conducted for violations of sphericity. Any significant interaction surviving FDR multiple-comparison adjustments were investigated via a one-way repeated measures ANOVA to determine which ROIs had differential activation for the various behaviors, and subsequent pairwise t-testing to determine how the *β*-coefficients for the behaviors differed within the ROI.

In preparation for exploratory group-level analyses, a gray matter mask was constructed in template space using Atropos priors (Tustison et al., 2014), which was multiplied by the intersection mask to produce a gray matter, intersection inclusion mask. Exploratory group-level analyses were then conducted on all voxels within this inclusion mask for each phase of the experiment using the Equitable Thresholding and Clustering method (ETAC; Cox, 2019) using the p-values 0.01, 0.005, and 0.001, blurs of 4, 6, and 8mm, nearest neighbors = 1, and two-sided thresholding. Any surviving clusters were used to extract mean *β*-coefficients from participant data. All scripts used in this experiment can be found at the experiment’s GitHub repository: https://github.com/nmuncy/STT.

## 3 Results

### 3.1 Behavioral Analyses

Analyses of Test1 performance yield a mean 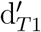 score of 0.66, indicating that successfully distinguishing between Targets and Lures was difficult, as the design of the experiment intended. Participants, however, were successful at detecting Targets and Lures in Test1, as the 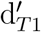 differed significantly from zero (*t*_(34)_ = 11.3, *q* < .001, 95% CI[.54, .78]). Response proportions were tested against chance (0.5) for each stimulus type; participants were more likely to Hit than Miss on Targets (*est* = .79, *χ*^2^ = 32.18, *q* < .0001, 95% CI[.69, .86]) while being equally likely to CR and FA on Lures (*est* = .43, *χ*^2^ = 1.43, *q* = .41, 95% CI[.33, .54]; Figure 2, left). On average, participants had 76 Hits, 54 FAs, 42 CRs, 20 Misses, and 6 non-responses in this phase. The 54 FA responses are of particular interest to the study: we designed the difficulty of the experiment to elicit approximately 60 FA responses in Test1 as two responses (Hit, FA) were possible in Test2, so subdividing Test1 FAs bins according to subsequent Test2 responses would yield sufficient trials to test statistically and model the hemodynamic response function.

**Figure 2:**
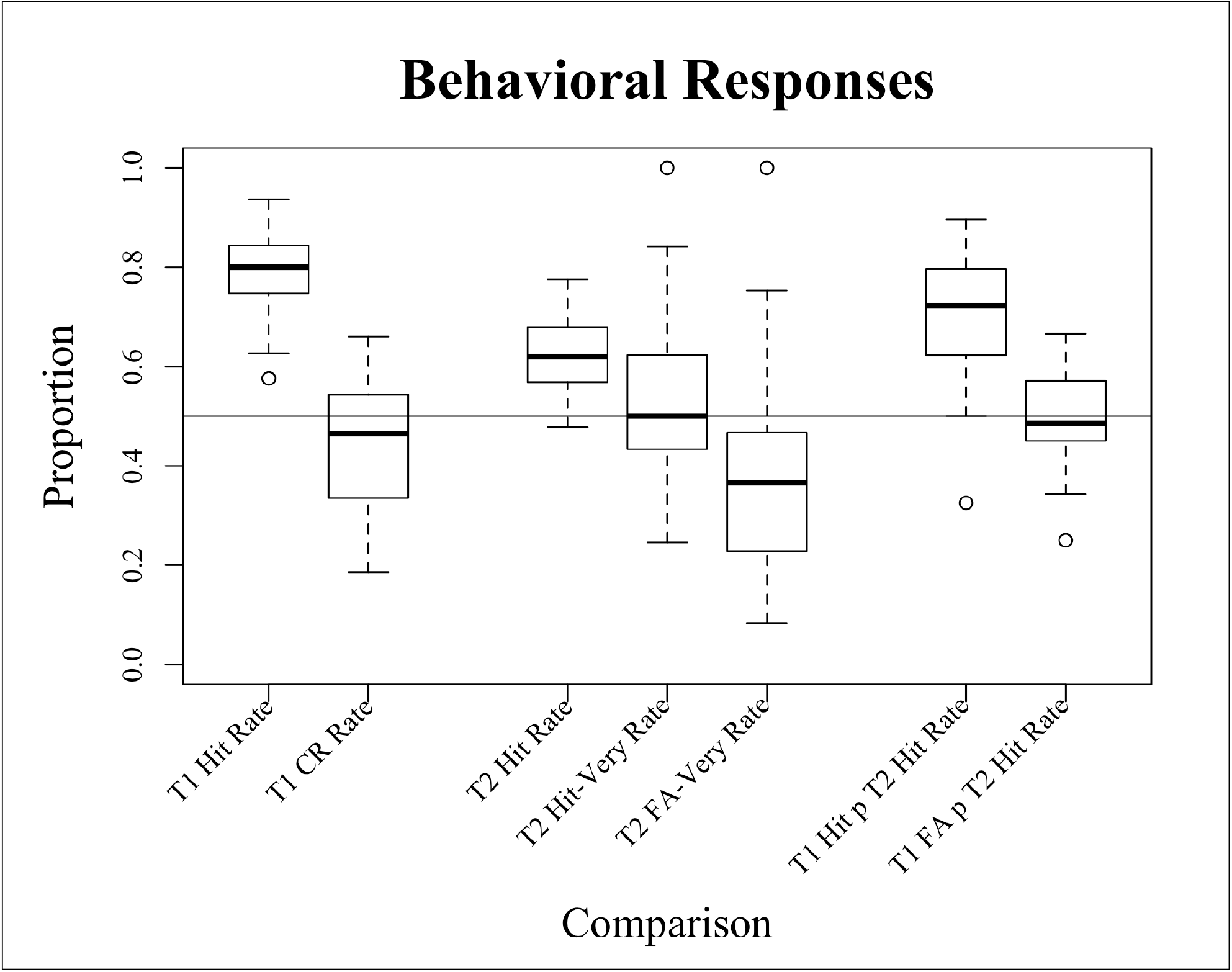
Boxplots of behavioral response proportions, for Test1 (T1), Test2 (T2), and Test1 preceding Test2 (T1 p T2). Performance at chance results in a proportion of 0.5. “Hit-Very Rate” = likelihood of responding Very Confident versus Somewhat Confident during Hits. “T1 Hit p T2 Hit Rate” = proportion of Test1 Hits that precede a Test2 Hit versus Test2 FA. Outliers are included.

Analyses of Test2 performance yielded a 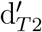 of 0.45, which also differed significantly from zero (*t*_(34)_ = 9.12, *q* < .0001, 95% CI[.35, .55]), and indicated that participants were more likely to Hit than FA (*est* = .62, *χ*^2^ = 11.43, *q* = .002, 95% CI[.55, .60]; Figure 2, middle). In terms of confidence, participants were equally likely to indicate Very and Somewhat confident during Test2 Hits (*est* = .53, *χ*^2^ = 11.43, *q* = .2, 95% CI[.43, .62]) and during Test2 FAs (*est* = .38, *χ*^2^ = 4.1, *q* = .09, 95% CI[.27, .49]). Finally, testing the d’ scores for Test1 versus Test2 indicated that participants performed better during Test1 (*t*_(34)_ = 4.37, *q* = .001, 95% CI[.11, .31]). On average, participants had 121 Hits, 73 FAs, and 5 non-responses.

Proportion testing of Test1 Hits and FAs which preceded Test2 responses revealed that a Test1 Hit was more likely to result in a subsequent Test2 Hit than a FA (*est* = .72, *χ*^2^ = 13.8, *q* < .001, 95% CI[.60, .82]), and interestingly that Test1 FAs were equally likely to result in either a subsequent Test2 Hit or FA (*est* = .5, *χ*^2^ ≈ 13.8, *q* ≈ 1, 95% CI[.37, .64]; Figure 2, right). These results, when viewed with respect to the Test2 results (i.e. a d’ which differs from 0 and a greater proportion of target detection), suggest that recovery from a Test1 FA is possible but not guaranteed. Analyses of the BOLD response during Test1 FAs that precede Hits versus FAs in the Test2 phase will help determine why some Test1 FAs are perpetuated into Test2 while others are corrected. Finally, it was possible that participants were simply selecting the Test2 item most recently encountered rather than the remembered Target from the Study phase. To this end we tested whether participants were more likely to select the Lure in Test2 following a Test1 CR response using proportion testing; we reasoned that a Test1 CR would result in a dissociable trace from the original, and that if a participant was simply selecting the most recently encountered stimulus in Test2 then they would select the Lure rather than the Target for these specific stimuli. Proportion testing determined that this was not the case (*est* = .33, *χ*^2^ = 3.57, *p* = .058, 95% CI[.20, .51]).

Test1 responses that preceded Test2 confidence responses consist of permutations of Test1 behaviors (Hit, Miss, CR, FA), Test2 behaviors (Hit, FA), and Test2 confidence responses (Very Confident, Somewhat Confident). It was originally thought that Test1_FA_-Test2_Hit_ responses would yield higher confidence responses than Test1_FA_-Test2_FA_, which could have lead to an interesting investigation of the BOLD response. Paired *t*-testing revealed, however, an equal number of “Very Confident” responses for both Hits and FAs in Test2 following a Test1 FA (*t*_(34)_ = 0.1, *q* = .9, 95% CI[*−*2.3, 2.5]), and indeed participants were equally likely to select “Very Confident” or “Somewhat Confident” during Test2 Hits and FAs following Test1 FAs according to proportion testing (*est* = .4, *χ*^2^ = 0.7, *p_F DR_* = .53, 95% CI[.23, .61]; *est* = .4, *χ*^2^ = 0.8, *q* = .53, 95% CI[.22, .59], respectively). Finally, parsing the data in such a fashion yielded bins that were too small to reasonably model the hemodynamic response (e.g. Test1 FA preceding Test2 Hit-Very = 11*±*6 responses on average). As such, all subsequent analyses were collapsed across confidence ratings.

Together, these behavioral findings indicate that a Test1 FA does not necessarily overwrite the original memory trace, as such a process would (a) suppress 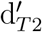 scores and (b) result in Test1 FA responses being more likely to precede a Test2 FA (indicated by a response proportion that approaches zero). Accordingly, these results support hypothesis (a) where we predicted that multiple representations could occur.

### 3.2 FMRI: Study trials preceding Test1 FAs preceding Test2 responses

Below we discuss the BOLD signal associated with overcoming mnemonic generalization during various aspects of the experiment. Specifically, we investigate processes during each phase of the experiment (Study, Test1, Test2) for their potential impact on overcoming a Test1 FA response. Analyses not involved with an attempted recovery from mnemonic generalization are reported in the Supplementary Material section as they do not specifically test the hypotheses of this experiment, e.g. a description of the BOLD during the Test1 phase of the experiment. In each section reported below, a series of analyses were conducted. First, *a priori* analyses utilizing hippocampal subregion masks are used to test hypothesis (b). Second, *a priori* analyses utilizing NeuroSynth ROI masks are used to test hypothesis (c). Finally, we note that (a) the paradigm employed in this experiment may differ substantially when compared to those from which the NeuroSynth ROI masks were derived, (b) that utilizing a single *β*-coefficient for an entire ROI is an insensitive approach, and (c) that simply creating ROI masks for all conceivable higher-order processes that could potentially be associated with the task would essentially result in testing the entire cerebrum. Accordingly, our third set of analyses utilize an exploratory *a posteriori* whole-brain approach in order to potentially identify any clusters that may be differentially involved in overcoming mnemonic generalization while accounting for spurious findings using an updated approach (Cox, 2019). This was done in order to test hypothesis (d). Also, note that the FDR method of correcting multiple comparisons was employed to account for the large number of analyses conducted in this experiment.

In order to investigate the effect that initial encoding may have in overcoming mnemonic generalization, trials from the Study phase of the experiment that preceded a Test1 FA response were coded for whether the behavior was corrected in Test2 (Hit) or not (FA; Supplemental Figure S4, purple). On average, there were 26 trials per behavioral bin, and one participant was excluded from this phase of the experiment due to excessive movement (n=34). Individual two-factor repeated measures ANOVAs, using ROI and Behavior as within-subject factors and the corresponding *β*-coefficient as the dependent variable, were used to investigate activity associated with the behavior within hippocampal subregions (left and right CA2/3/DG and CA1) as well as “Memory Encoding” NeuroSynth ROIs. Following a correction for sphericity, no ROI × Behavior interaction was detected in this phase of the experiment within the hippocampal subregions (*F*_(3,99)_ = 2.58, 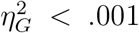, *GG_ε_* = .79, *p_GG_* = .07) nor was an interaction detected within the NeuroSynth ROIs (*F*_(3,99)_ = 2.77, 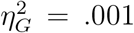, *GG_ε_* = .58, *p_GG_* = .08). Further, exploratory whole-brain analyses failed to detect any regions during the encoding phase that discriminated for whether a subsequent Test1 FA would be corrected or perpetuated in Test2. As such, we fail to demonstrate any evidence that encoding processes play a role in whether a Test1 FA will be corrected and therefore fail to gather any evidence for hypotheses (b-d) from this phase of the experiment.

### 3.3 FMRI: Test1 preceding Test2

Test1 trials which resulted in FA responses were coded according to whether or not the response would be corrected in Test2 (Supplemental Figure S4, red). This was done in order to investigate any potential processes occurring during the Test1 FA that may allow for subsequent recovery from mnemonic generalization. Two participants were excluded from these analyses due to excessive movement in the Test1 phase (n=33) and behavioral bin sizes were identical to those reported above. A set of two-factor repeated measures ANOVAs identical to those above were used investigate a potential ROI × Behavior interaction. As above, such analyses failed to detect a significant interaction within the hippocampal subregions CA2/3/DG or CA1 (*F*_(3,96)_ = 1.39, 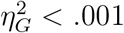, *p* = .24) as well as among NeuroSynth”Memory Retrieval” ROIs (*F*_(9,288)_ = 0.65, 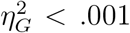, *p* = .74). Separately, it is possible that whether a Test1 FA will be corrected in Test2 is dependent on the encoding quality of the Test1 Lure in addition to retrieval processes. As such, we conducted a similar two-factor repeated measures ANOVA using the NeuroSynth regions associated with “Memory Encoding” but again failed to detect a significant interaction (*F*_(3,96)_ = 1.56, 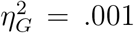, *p* = .2).

Exploratory analyses were conducted in order to investigate whether higher-order processes were associated with perpetuating or correcting a FA. While NeuroSynth queries for terms such as “Decision Making”, “Attention”, “Executive Function”, etc. could have been performed, such an approach would have increased the already large number of statistical tests conducted in this experiment, an issue that is additional to those raised in Section 3.2. Accordingly, we tested hypothesis (d) via an exploratory whole-brain analysis. Such an analysis of Test1 FA trials which preceded a Test2 Hit (FpH) versus FA (FpF) implicated two regions as being differentially engaged: the left insular cortex (LINS) and right supramarginal gyrus (RSMG; Figure 3). Both regions were more active during a Test1 FA preceding a Test2 Hit than a Test2 FA (Table 1). Such a pattern is consistent with increased activity in attention networks during Test1 FAs being predictive of a subsequent correction of the memory mistake (see Discussion).

**Figure 3:**
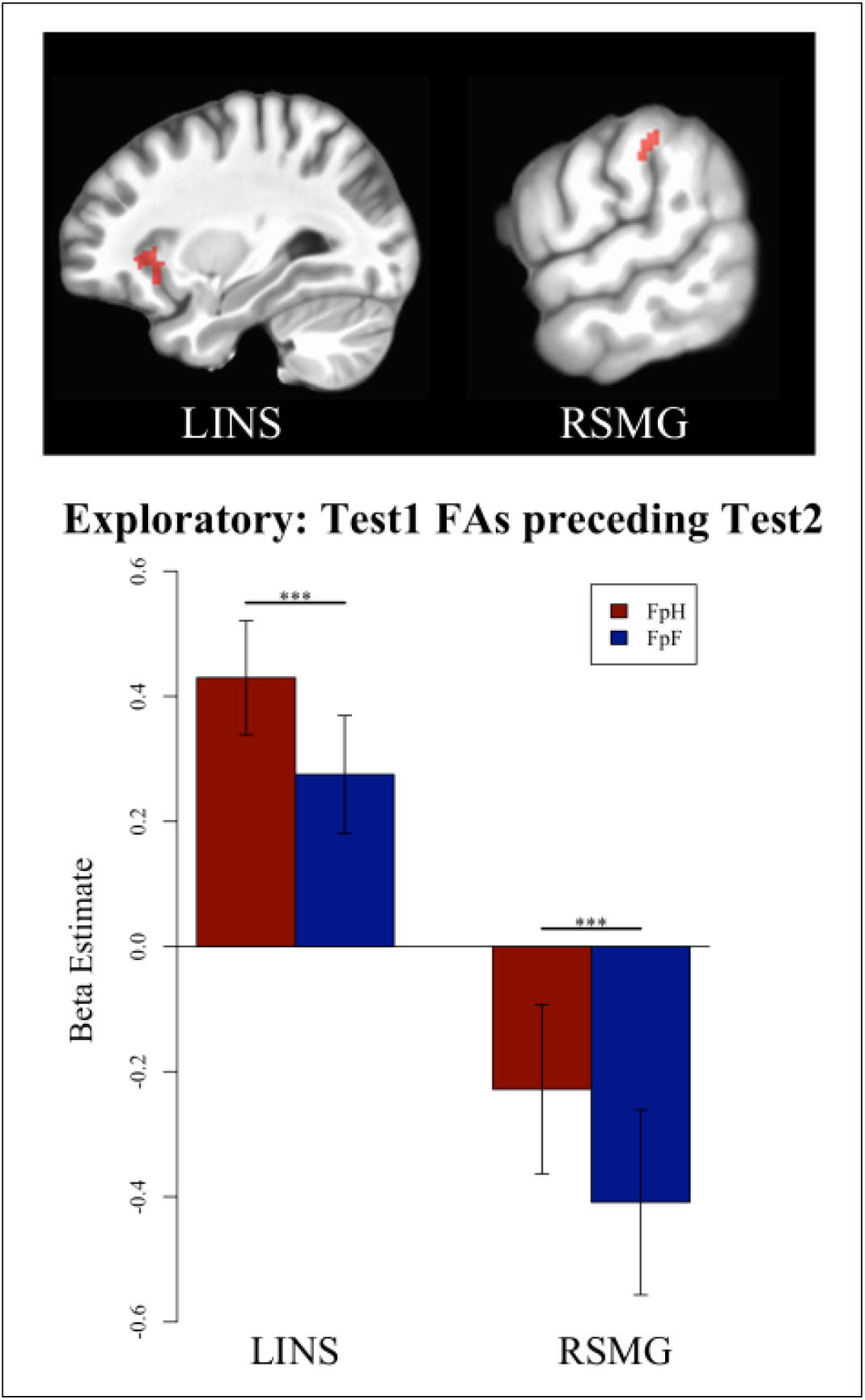
Clusters which demonstrate differential signal during Test1 FAs that precede a Test2 Hit versus FA. LINS = left insular cortex, RSMG = right supramarginal gyrus. FpH = Test1 FA preceding a Test2 Hit, FpF = Test1 FA preceding a Test2 FA.

**Table 1:** Cluster sizes and test statistics for the exploratory whole-brain analyses of Test1 FA trials preceding Test2. ROI = region of interest, K = cluster size, X-Z = coordinate (peak), T(xx) = t-test(df), Sig = significant, *** = p<.0001, LB = 95% confidence interval lower bound, UB = upper bound. LINS = left insular cortex, RSMG = right supramarginal gyrus.

| ROI | K | X | Y | Z | T(32) | Sig | LB | UB |
| --- | --- | --- | --- | --- | --- | --- | --- | --- |
| LINS | 98 | 31 | -26 | -3 | 4.7 | *** | 0.05 | 0.12 |
| RSMG | 19 | -63 | 30 | 41 | 6.8 | *** | 0.08 | 0.15 |

With respect to the *a priori* and *a posteriori* findings above, then, we do not detect any evidence in support of hypothesis (b) in that no significant differences were found within hippocampal subregions, nor for hypothesis (c). We do, however, find support for hypothesis (d), as regions associated with higher-order processes (e.g. attention networks) were differentially engaged during the Test1 FA according to whether or not the behavior would subsequently be corrected.

### 3.4 FMRI: Test2 responses following Test1 FAs

Test2 responses (Hit, FA) following a Test1 FA were coded in order to investigate any potential processes occurring during the second testing phase involved in overcoming interference resulting from a Test1 FA (Supplemental Figure S4, green). Three participants were excluded from analyses of this last phase due to excessive movement during Test2 (n=32), and again, there were 26 trials per bin on average. As above, a two-factor repeated measures ANOVA failed to detect a significant interaction within either the hippocampal subregions (*F*_(3,93)_ = 1.1, 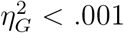, *p* = .43) or the NeuroSynth “Memory Retrieval” ROIs (*F*_(9,279)_ = 0.64, 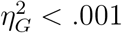, *p* = .75). Finally, exploratory whole-brain analyses did not detect any regions that differed significantly between their BOLD signal associated with Test2 Hits versus FAs following a Test1 FA. To summarize, we did not observe any evidence from the Test2 phase in support of hypotheses (b-d).

## 4 Discussion

This experiment set out to investigate the impact of a false memory response on the original memory representation, and to determine what, if any, neural processes were associated with recovering the original memory trace. To this end, we employed a Study-Test1-Test2 paradigm within an MRI scanner, in which participants were trained on a series of images (Study), tasked with identifying either Targets or similar Lures presented individually (Test1), and then tasked with identifying the Target when both Targets and Lures were presented simultaneously (Test2). False Alarm (FA) responses in Test1 were operationalized as false memories and were used to investigate whether participants could correctly identify the Target during Test2 (Hit) following a Test1 FA. Briefly, we hypothesized (a) that a FA would write a memory trace of the Lure without necessarily overwriting the Target’s trace such that the two representations would be dissociable, and (b) that the CA2/3/DG regions of the hippocampus as well as (c) encoding and retrieval nodes and (d) higher-order executive processes would be integral to the formation of a second, dissociable representation.

As anticipated in the paradigm design, participants made a large number of FA responses in Test1, and proportion testing revealed that participants were equally likely to correct (Hit) and perpetuate (FA) the mistake (Test1 FA) into Test2 (Figure 2). That is, a FA does not unambiguously lead to overwriting the Target memory trace, as would be indicated by a greater likelihood to perpetuate the FA response in Test2, but neither does the FA necessarily lead to writing multiple representations from which the Target is always dissociable. Given that participants were indeed successful in Test2 (Section 3.1), the behavioral data are ambiguous as to the effect of the FA on the Target trace, and so from the behavioral analyses alone we are unable to reject the null of hypothesis (a). We therefore turn to analyses of the corresponding BOLD signal to potentially shed some light on why some FA responses are corrected and others perpetuated.

Three sets of functional analyses investigated the various phases for the experiment, each set containing *a priori* analyses on hippocampal subregion and NeuroSynth masks and *a posteriori* whole-brain exploratory analyses. First, we investigated potential processes occurring during mnemonic generalization that might have predicted whether the original trace was recovered in a subsequent test phase, i.e., we investigated whether differential BOLD signal existed during Test1 FA responses that predicted the Test2 fate (Hit versus FA). While no differential signal was detected in the *a priori* hippocampal subregion, NeuroSynth “Memory Encoding”, and NeuroSynth “Memory Retrieval”, the exploratory whole-brain analysis revealed two clusters (see Section 3.3). The left insular cortex and right supramarginal gyrus were both detected to be more active during FAs preceding a Test2 Hit relative to those preceding a Test2 FA response. As these regions have previously been associated with saliency processing and bottom-up attention networks (Cabeza et al., 2008; Menon & Uddin, 2010; Seeley et al., 2007), one possible interpretation, then, is that increased activity in these regions during a Test1 FA may be associated with better attention to the relevant details of the stimulus which, while sub-threshold for successful Lure detection, nevertheless led to the formation of a Lure representation such that the Target and Lure memory traces were subsequently dissociable in Test2.

Second, we investigated BOLD signal from the Study phase to determine if encoding had a differential impact on whether the memory trace would be recoverable following a memory mistake (see Section 3.2). We did not detect any differential signal that predicted whether a Test1 memory mistake (FA) would be corrected in Test2 (Hit) in the hippocampal subregions, NeuroSynth “Memory Encoding” regions, or with an exploratory analysis. This was somewhat surprising as we expected to find evidence indicating a range of encoding efficiencies preceding differing Test2 fates for stimuli that elicited Test1 FA responses. Instead, we did not detect any evidence that initial encoding processes played a role in whether the original memory trace was recoverable following mnemonic generalization. These null results are in contrast to those resulting from investigating whether processes during encoding differentially predicted Test1 performance (see Supplementary Material). In that set of analyses, multiple regions were differentially engaged, regions which were consistent with the subsequent memory and forgetting literature (Cabeza et al., 2008; H. Kim, 2011; Shrager et al., 2008). Specifically, differential BOLD signal was detected during the encoding phase in numerous regions associated with recognition memory, attention, and the default mode network that preceded Test1 performance. It is interesting, then, that while signal was detected during the Study phase that is consistent with established work on subsequent memory and forgetting, we did not detect any signal during this phase that predicted the fate of subsequently generalized items. Perhaps it is possible that initial encoding processes do not have a role in overcoming subsequent mnemonic generalization, but this seems to be an over interpretation of null findings (Chen et al., 2017) and it is more likely that our given contrasts and/or analyses did not have the requisite sensitivity. Indeed, collapsing across entire ROIs will average potentially relevant signal with irrelevant signal, deflating test statistics. And while the ETAC (Cox, 2019) approach is designed to be sensitive to both small, clusters with large test statistics as well as larger clusters with lower statistics, perhaps the critical region and signal is too small and similar to discriminate with the methods employed in this analysis.

Third, we investigated potential processes involved in overcoming mnemonic generalization by assessing whether differential processes in Test2 were associated with Hits versus FAs following a Test1 FA (see Section 3.4). We reasoned that, when confronted with both the Target and Lure during Test2, participants might recognize their previous mistake (Test1 FA) and then correctly select the Target. Surprisingly, again, we did not detect any differential signal in any of the analyses during this phase of the experiment associated with overcoming mnemonic generalization. We hypothesize that these null results are, as above, due to a lack of sensitivity in techniques or measures and that processes involved in overcoming interference would facilitate retrieval of a memory trace following mnemonic generalization.

Some results of this study were surprising. First, participants were equally likely to Hit or FA in Test2 to stimuli that had previously resulted in a Test1 FA (Section 3.1, Figure 2, right). We hypothesized that a FA would produce a second, dissociable memory trace and so we expected to see a bias towards Target selection in Test2 for stimuli that previously elicited a Test1 FA. We also expected that if our hypothesis was incorrect (and a mnemonic discrimination overwrites the original trace), then we would detect a Test2 bias towards the Lure. An ambiguous result, then, does not offer compelling evidence for or against the hypothesis. It is possible that this result is a function of the experimental design itself: a large number of difficult Target-Lure pairs were utilized in order to elicit behavior performance such that dividing the number of Test1 FA responses according to subsequent Test2 behaviors would yield an appropriate number trials for modeling the hemodynamic response function. To this end, the task was successful in that it elicited approximately 60 Test1 FA responses and a corresponding 30 Test2 Hits and FAs following such Test1 FAs from each participant, on average. Unfortunately, however, this design resulted in a task that lasted 1.5 hours. Despite such a long, arduous task, evidence suggest that participants remained engaged in the task: the Test2 d’ differed significantly from chance, and proportion testing reject the notion that participants were simply selecting the Test2 stimulus they had most recently encountered rather than attempting to select the Target presented in the Study phase. We propose, then, that participants were equally likely to select a Test2 Hit or FA following a Test1 FA due to significant amounts of interference and fatigue. We also find it likely that a mnemonic generalization does in some instances overwrite the original trace, especially given that differential processes were detected in Section 3.3, and the fact that only some memory traces are overwritten would produce such results. We note that the response proportion of Test1 FA preceding a Test2 Hit, while at chance (see Figure 2, Right), in no way approached 0, a score which would indicate a greater tendency for the mnemonic generalization to overwrite the original representation.

A second unexpected result was the lack of findings within the hippocampal subregions in all analyses save those of Test1 (see Supplementary Materials). While we demonstrate the differential roles of left CA1 and CA2/CA3/DG in this task with respect to Hits and CRs, in which the hippocampus appeared to be biased towards Target detection in this task, the lack of subregion findings in other analyses of the task was surprising to us given the wealth of literature that has well established the roles of these regions in similar tasks (Bakker et al., 2008; J. Kim & Yassa, 2013; Kirwan & Stark, 2007; Lacy et al., 2011; Rolls & Kesner, 2016). Accordingly, we cannot reject the null of hypothesis (b) as we did not detect differential subregion processing during the FA that predicted subsequent performance. These null results are perhaps a type II or false negative error which resulted from extracting a single parameter estimate for the entire ROI, thereby collapsing across a significant amount of noise as well as signal, possibly washing out the effect. Indeed, it has been noted that the long-axis of the hippocampus contains a functional gradient and discrete domains (Strange et al., 2014), so collapsing across all variance along the anterior-posterior axis of the hippocampal subregions may be inappropriate. Similarly, our contrast of Test1 FA signal coded for subsequent Test2 performance may have simply failed to detect a difference between those two conditions in terms of subregion BOLD signal, but this does not necessarily imply that the hippocampus is not differentially processing the lures such that subsequent behavior results in a Hit versus FA. Rather, the processes involved in such a computation could be rather similar to one another and so a comparison of the mean subregion BOLD signal associated with each process may again be simply too insensitive.

Third, we were surprised by the lack of significant differences for the fMRI data of the Test2 phase (see, Supplementary Materials). The behavioral data indicate that successful Target detection was possible, and that participants were more likely to Hit in Test2 following a Test1 Hit, so performance was not at chance. The lack of findings may not be unwarranted, however, as there are very few studies reporting positive results from two-alternative, forced choice, recognition memory paradigms using fMRI. One notable exception by Olsen and colleagues (2009) is somewhat similar to this experiment in terms of participants and power. Olsen et al. (2009) demonstrated that the medial temporal region was active while maintaining the memory of a set of faces during a delay period prior to a two-alternative forced choice test, an area similar to where we expected activation to occur. The lack of findings in the present experiment may be due to differences between the experiments in fMRI processing methods. One stark difference is in the methods modeling both the noise and auto-correlation function at the group-analysis step employed in this experiment, as this improvement is more conservative than previous random-field based models in terms of false discovery rates but was only recently developed (Cox, 2018, 2019; Eklund et al., 2016). Furthermore, while significant differences were not detected within the fMRI data at the group level, one should not assume that “[i]f the result is not statistically significant, then it proves that no effect or difference exists” (Chen et al., 2017, p. 955). Indeed, as Chen et al. 2017 points out, null results may very well be the result of sub-threshold signal and/or a lack of differential signal between our chosen contrasts at our resolution.

In conclusion, the evidence presented here suggest that mnemonic generalization processes do not necessarily overwrite the original memory trace, but that the original memory trace is recoverable following failed discrimination in some instances. These data also suggest that increased activity in the left insular cortex and right supramarginal gyrus during mnemonic generalization is predictive of whether the original trace will be recovered, findings which may implicate saliency and attention networks. With respect to our hypotheses, we find some support for hypotheses (a, d), i.e. reject their null, but we gathered no evidence to support rejecting the null of hypotheses (b, c).

## 5 Supplementary Material

### 5.1 Supplementary Methods

For *a priori* analyses, mean *β*-coefficients were extracted from regions of interest (ROIs) within hippocampal subregions (Supplemental Figure S1), and NeuroSynth regions associated with the search terms “Memory Encoding” (Supplemental Figure S2) and “Memory Retrieval” (Supplemental Figure S3), also Supplemental Table S1. All masks were down-sampled into functional dimensions and exist in MNI space.

**Figure S1:**
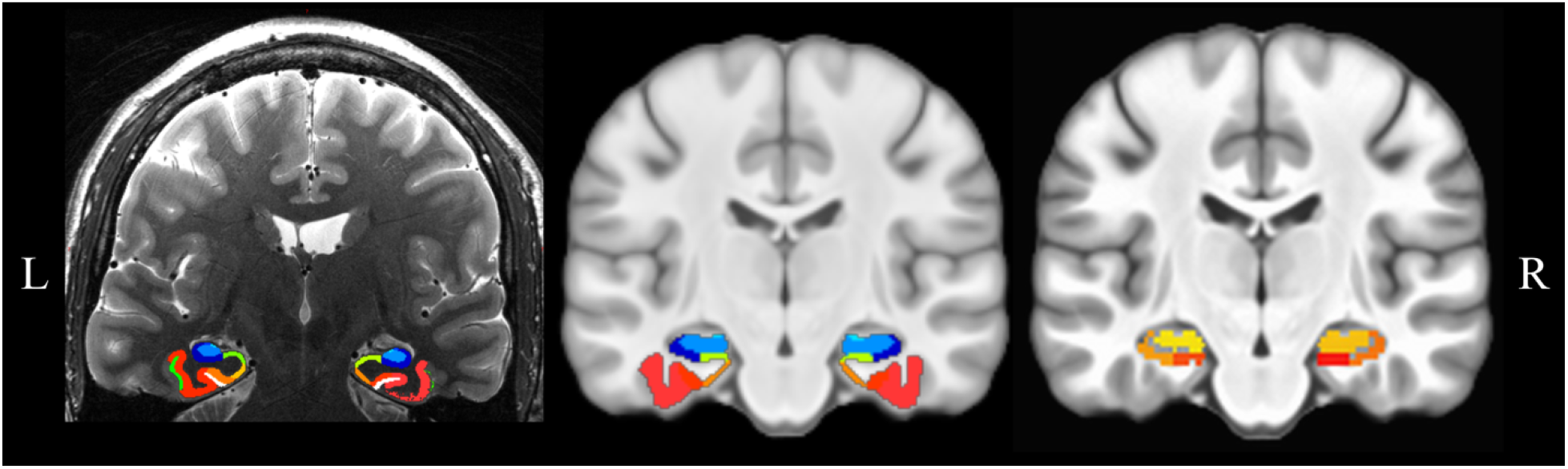
Left, a participant’s T2-weighted scan segmented for the medial temporal lobe via ASHS. Middle, template medial temporal lobe masks constructed in MNI space via Joint Label Fusion. Right, resampled masks, where overlapping labels were excluded.

### 5.2 Supplementary Results

#### 5.2.1 Study trials preceding Test1 responses (SpT1)

Study stimulus responses were coded for subsequent Test1 behavior in order to investigate the potential effect of encoding on Test1 performance. For example, one stimulus in Study could elicit a Hit in Test1 while a different stimulus could elicit a FA response (Supplemental Figure S4, blue). Separate *a priori* analyses investigated hippocampal sub-regions (CA1, CA2/3/DG) and NeuroSynth “Memory Encoding” regions of interest using two-factor within-subject ANOVAs (ROI × Behavior) using the mean *β*-coefficient as the dependent variable; performance on Lures (i.e. encoding preceding CR or FA) and Targets (i.e. encoding preceding Hit or Miss) were investigated separately. An ROI × Behavior interaction was not detected among hippocampal subregions preceding Lure or Target performance (*F*_(3,99)_ = 1.28, 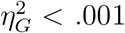, *p* = .2; *F*_(3,99)_ = 0.06, 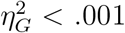, *p* = .9, respectively), nor was such an interaction was detected among NeuroSynth encoding ROIs preceding Lure or Target performance (*F*_(3, 99)_ = 0.37, 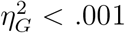, *p* = .7; *F*_(3,99)_ = 2.19, 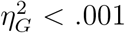, *p* = .09, respectively).

**Figure S2:**
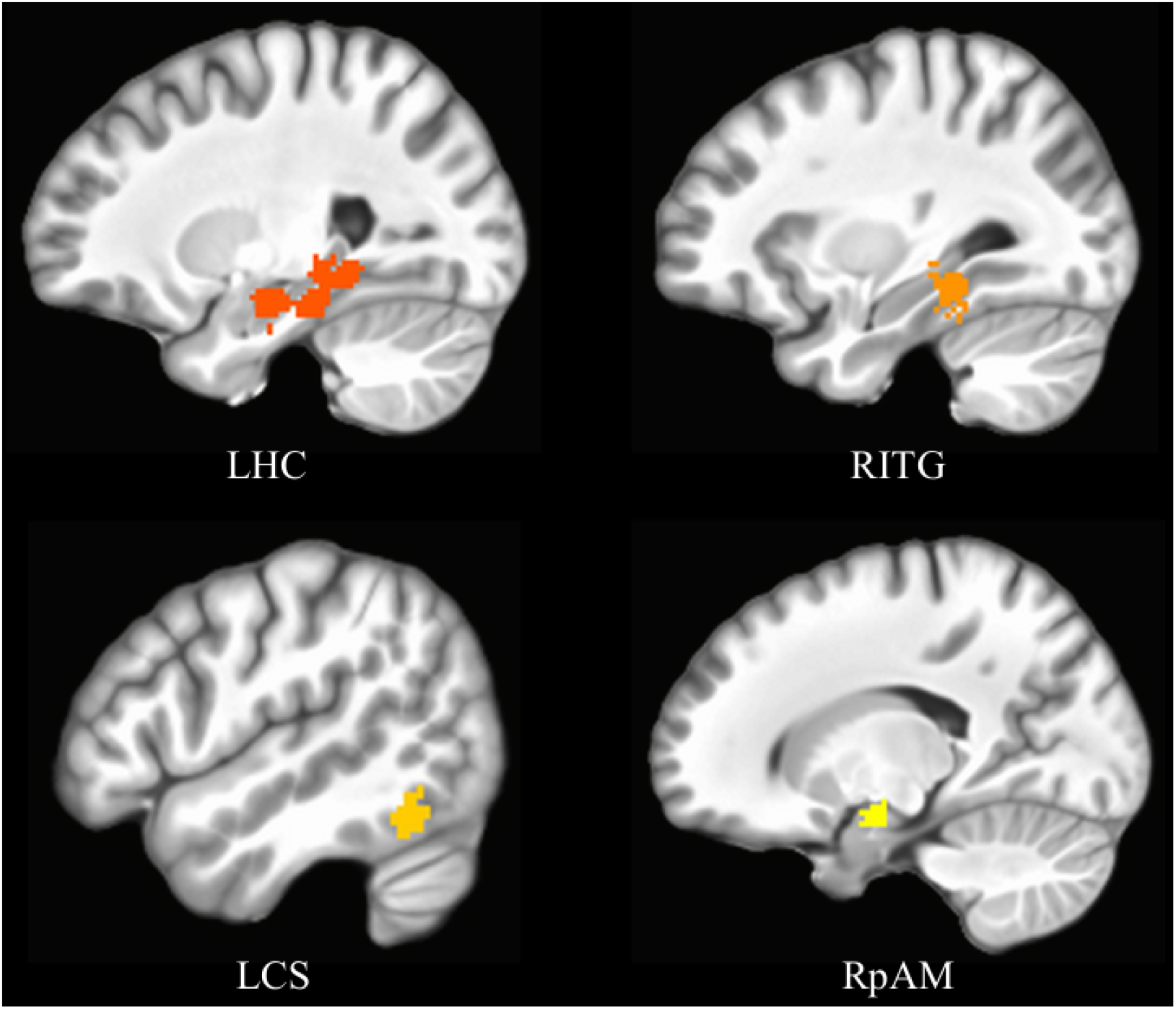
NeuroSynth masks associated with “Memory Encoding”. See Supplemental Table S1 for abbreviations.

#### 5.2.2 Test1 responses

Test1 trials were coded for the relative success of target and lure detection, using the labels Hit, FA, CR, and Miss. *A priori* analyses were identical to those in Section 5.2.1 save that the NeuroSynth ROIs were derived from the search term “Memory Retrieval” and the *β*-coefficients were extracted from the Test1 phase of the experiment. For the hippocampal subregions, no significant interaction between the factors were detected during Lure trials (*F*_(3,96)_ = 1.15, 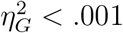, *p* = .3), nor was an interaction detected for Target trials following a correction for sphericity (*F*_(3,96)_ = 2.91, 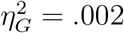, *GG_E_* = 0.65, *p_GG_* = .06).

**Figure S3:**
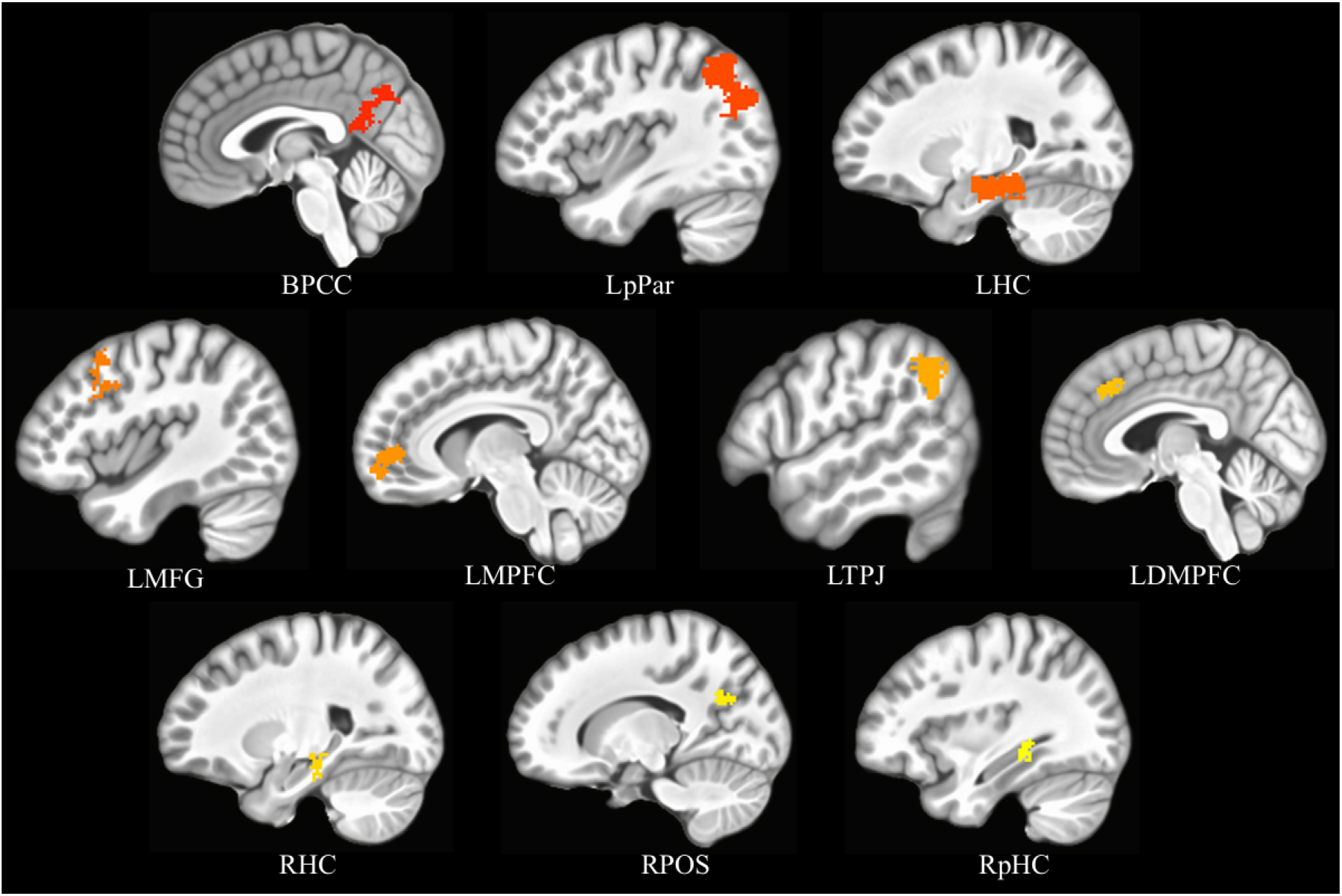
NeuroSynth masks associated with “Memory Retrieval”. See Supplemental Table S1 for abbreviations.

**Table S1:** Description of NeuroSynth masks. Group = search term, ROI = region of interest, Abbv = abbreviation, X-Z = center of mass for MNI coordinate X-Z.

| Group | ROI | Abbv | Vol | X | Y | Z |
| --- | --- | --- | --- | --- | --- | --- |
| Encoding | L. hippocampus | LHC | 776 | -26 | -24 | -14 |
|  | R. inferotemporal gyrus | RITG | 217 | 31 | -34 | -14 |
|  | R. collateral sulcus | LCS | 89 | -51 | -55 | -16 |
|  | R. posterior amygdala | RpAM | 83 | 17 | -7 | -18 |
| Retrieval | B. posterior cingulate | BPCC | 1194 | -3 | -61 | 27 |
|  | L. posterior parietal | LPPar | 946 | -39 | -66 | 40 |
|  | L. hippocampus | LHC | 735 | -23 | -25 | -17 |
|  | L. middle frontal gyrus | LMFG | 394 | -42 | 17 | 44 |
|  | L. medial prefrontal | LMPFC | 253 | -8 | 56 | -1 |
|  | L. temporoparietal junction | LTPJ | 247 | -56 | -54 | 37 |
|  | L. dorsal medial prefrontal | LDMPFC | 149 | -4 | 32 | 38 |
|  | R. hippocampus | RHC | 146 | 24 | -25 | -14 |
|  | R. parieto-occipital sulcus | RPOS | 115 | 15 | -63 | 26 |
|  | R. posterior hippocampus | RpHC | 114 | 34 | -36 | -6 |

**Figure S4:**
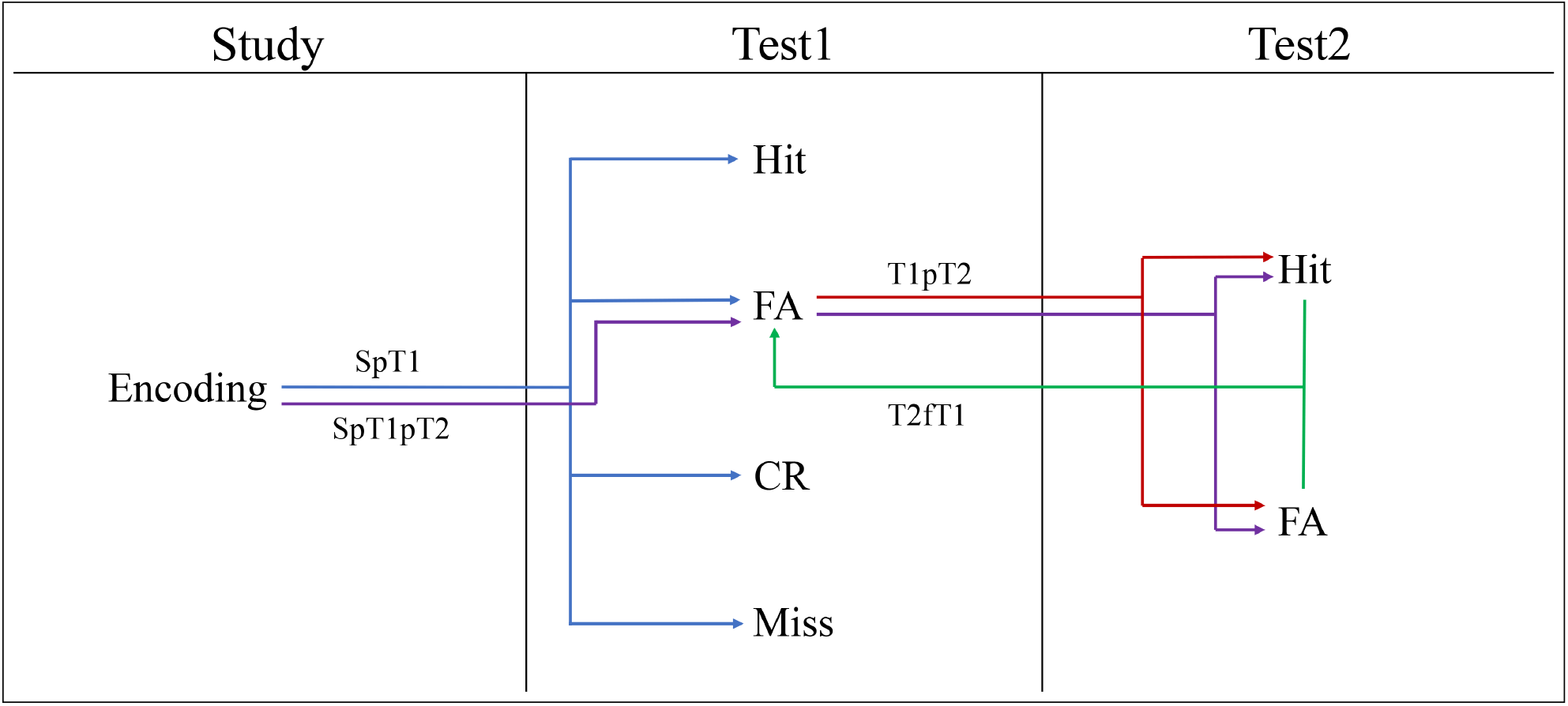
Schematic of analyses performed. The experiment consists of three phases, Study, Test1, Test2. Study trials were coded for which stimuli preceded Test1 responses (SpT1, blue) as well as the resulting response in Test2 following a Test1 response (SpT1pT2, purple). Test1 trials were analyzed both for the behavior which occurred in Test1 (not colored) as well as for responses which preceded a Test2 response (T1pT2, red). Similarly, Test2 trials were analyzed for Test2 behaviors (not colored) as well as for which Test1 behavior the Test2 response followed (T2fT1).

Conversely, NeuroSynth Retrieval ROIs were differentially engaged with both Lure and Target trials (*F*_(9, 288)_ = 10.6, 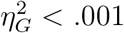, *GG_∊_* = 0.46, *p_GG_* < .001; *F*_(9,288)_ = 15.3, 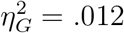, *GG_∊_* = 0.59, *p_GG_* < .001, respectively). *Post hoc* two-sided paired *t*-testing of the Lure trials revealed that three regions were driving the ROI × behavior interaction: the left middle frontal gyrus, left temporoparietal junction, and left dorsal medial prefrontal cortex. Each cluster was more active during CR relative to FA responses. Likewise, subsequent testing revealed that four clusters were driving the interaction for Target trials: the left dorsal medial prefrontal cortex, left middle frontal gyrus, left hippocampus, and left medial prefrontal cortex (Supplemental Figure S3, Supplemental Table S1). The former two clusters were more active during Miss trials, relative to Hit, and the latter two clusters had the opposite pattern of activity (Supplemental Table S2).

**Table S2:** NeuroSynth Retrieval statistics for Test1 analyses. LTPJ = left temporoparietal junction, LDMPFC = left dorsal medial prefrontal cortex, LMFG = left middle frontal gyrus, LHC = left hippocampus, LMPFC = left medial prefrontal cortex. ** = p<.01, *** = p<.001. LB = lower bound of 95% confidence interval, UB = upper bound of 95% confidence interval.

| Contrast | ROI | T(32) | Sig | LB | UB |
| --- | --- | --- | --- | --- | --- |
| CR vs FA | LTPJ | 3.6 | *** | 0.018 | 0.065 |
|  | LDMPFC | 4.7 | *** | 0.085 | 0.216 |
|  | LMFG | 3.3 | ** | 0.022 | 0.091 |
| Hit vs Miss | LDMPFC | -4.4 | *** | -0.215 | -0.0812 |
|  | LMFG | -3.8 | *** | -0.09 | -0.027 |
|  | LHC | 2.7 | ** | 0.012 | 0.085 |
|  | LMPFC | 6.1 | *** | 0.096 | 0.194 |

#### 5.2.3 Test2 Responses

A two-factor repeated measures MANOVA failed to detect a significant interaction of hippocampal subregions and Test2 behaviors (Hit, FA; *F* (5,155) = 1.04, 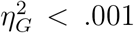, *GG_ε_* = .75, *p*(*GG*)_*FDR*_ = .39) or among NeuroSynth ROIs associated with retrieval (*F* (9,279) = .19, 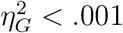, *GG_ε_* = .51, *p*(*GG*)_*FDR*_ = .99).

